# Elevated *P2×7r* and *P2×4r* transcripts levels in the Flinders Sensitive Line Rats, a genetic animal model of depression

**DOI:** 10.1101/395194

**Authors:** Deidiane Elisa Ribeiro, Heidi Kaastrup Müller, Betina Elfving, Samia Regiane Lourenço Joca, Gregers Wegener

## Abstract

P2×7 and P2×4 receptors (P2×7R and P2×4R, respectively) are ligand-gated ion channels activated by adenosine triphosphate (ATP), which have been associated to dysfunctional processes in stress responses linked to depression, such as neurotransmitter release, cognition, sleep, energy, appetite, immune and endocrine dysfunction. Clinical studies indicate that polymorphisms in the *P2×7r* gene results in increased susceptibility for development of depression. Existing studies have investigated the role of P2×7R and P2×4R in animal models based on stress exposure. Therefore, the present work aimed to investigate the transcript and protein levels of these receptors in a genetic animal model of depression, the Flinders Sensitive Line (FSL) and its control group, the Flinders Resistant Line (FRL) rats. We found that FSL rats have increased transcript levels of P2×7R and P2×4R in frontal cortex (FC), ventral and dorsal hippocampus (vHip and dHip, respectively) compared to FRL rats. There were no alterations in the protein levels in the FC and dHip, but the P2×7R was lower in FSL than in FRL rats in the vHip. The results suggest that increased transcripts levels of *P2×7r* and *P2×4r* in the FSL rats may contribute to the stress-susceptibility observed in these animals.

## 1 Introduction

Stress exposure is a key factor for development of depression (1) and both are associated with dysfunctions in common neuronal transmitters, circuitries and receptor systems (2). The P2×7 and P2×4 receptors (P2×7R and P2×4R) are ligand-gated ion channels activated by adenosine triphosphate (ATP) which have been indicated to modulate such process (3). Interestingly, existing studies support the importance of P2×7R in depression and stress response. Polymorphisms in the *P2×7r* are associated with increased susceptibility for development of depression (4, 5) and linked to symptom-severity (6). Genetic deletion of P2×7R results in a transgenic mouse with less depressant phenotype, i.e. decreased immobility time in the forced swim test (FST) and tail suspension test (TST) (7). These animals also showed attenuated anhedonia response in the sucrose preference test after bacterial endotoxin challenge (8).

Independently, our research group and Csolle et al. described for the first time that the treatment with P2XR antagonists induce antidepressant-like effect in the TST (9) and in the FST (10). Subesquent studies demonstrate that repeated administration of a P2×7R antagonist attenuated the elevated immobility time in the TST and FST induced by treatment with lipopolysaccharide (11), reversed anhedonia caused by chronic unpredictable stress (12), and helplessness behavior induced by inescapable foot shocks (13).

Similarly, the P2×4R also appears to be involved in response to stress. Acute treatment with a positive allosteric modulator of P2×4R, ivermectin, increased depressive-like behaviour in animals exposed to the FST and TST in low doses (14), but in higher doses induced antidepressant-like effect in the TST (14).

As most studies involving P2×7R and P2×4R link these receptors to stress response, the present study was conducted using the Flinders Sensitive Line (FSL) and the Flinders Resistant Line (FRL) rats, a genetic animal model of depression (15-17). Therefore, the work aimed to investigate the transcript and protein levels of P2×7R and P2×4R in frontal cortex (FC), ventral and dorsal hippocampus (vHip and dHip, respectively), which are key structures in stress and depression neurobiology, of FSL/FRL rats.

## 2 Materials and Methods

### 2.1 Animals

Male FSL/FRL rats (10 weeks old) obtained from the breeding colonies at the Translational Neuropsychiatry Unit, Aarhus University, Denmark, were kept in the vivarium, housed in pairs in Euro Standard Type III-H cages, under standard conditions: 22 ± 1°C, lights on from 6:00 a.m. to 6:00 p.m., free access to food and water.

### 2.2 Experimental design

FSL and FRL (control group) rats were decapitated without prior anesthesia, the brain was removed, FC, vHip and dHip were dissected, frozen on powdered dry-ice and maintained at −80°C until analysis. Transcripts levels of P2×7R and P2×4R were investigated by real-time quantitative polymerase chain reaction (qPCR) while protein expression was analysed by Western blotting (WB).

### 2.3 RNA and protein isolation

RNA and protein were isolated from the samples of frontal cortex, ventral and dorsal hippocampus with PARIS™ kit (Ambion, TX, USA), according to the manufacturer’s specifications and previous studies (18, 19).

#### 2.3.1 Real-time qPCR

Transcript levels were determined in samples of frontal cortex, ventral and dorsal hippocampus from FSL/FRL rats with real-time qPCR, as described before (18, 19).

The RNA concentration and the purity were determined by a NanoDrop 1000 spectrophotometer (Thermo Fischer Scientific). Before cDNA synthesis, the RNA concentration of the samples was adjusted to match the sample with the lowest concentration (111.08 ng/μL). RNA was reversely transcribed using random primers and Superscript IV Reverse Transcriptase (Invitrogen, CA) following manufacturer’s instructions. The cDNA samples were stored undiluted (61.09 ng/μL) at −80 °C until real-time qPCR analysis.

The samples were diluted with DEPC water prior to real-time qPCR analysis (1:40). The real-time qPCR reactions were carried out in 96-well PCR-plates using an Mx3005P (Stratagene, USA) and SYBR Green.

The expression of eight different reference genes (*18sRNA, ActB, CycA, Gapd, Hmbs, Hprt1, Rpl13A, Ywhaz*) and 2 target genes (*P2×7r* and *P2×4r*) were investigated. The reference genes were selected as described in (20). Essential gene data about primer sequence and amplicon sizes are given in Table 1. The primers were obtained from Sigma-Aldrich, Denmark.

**Table 1.**
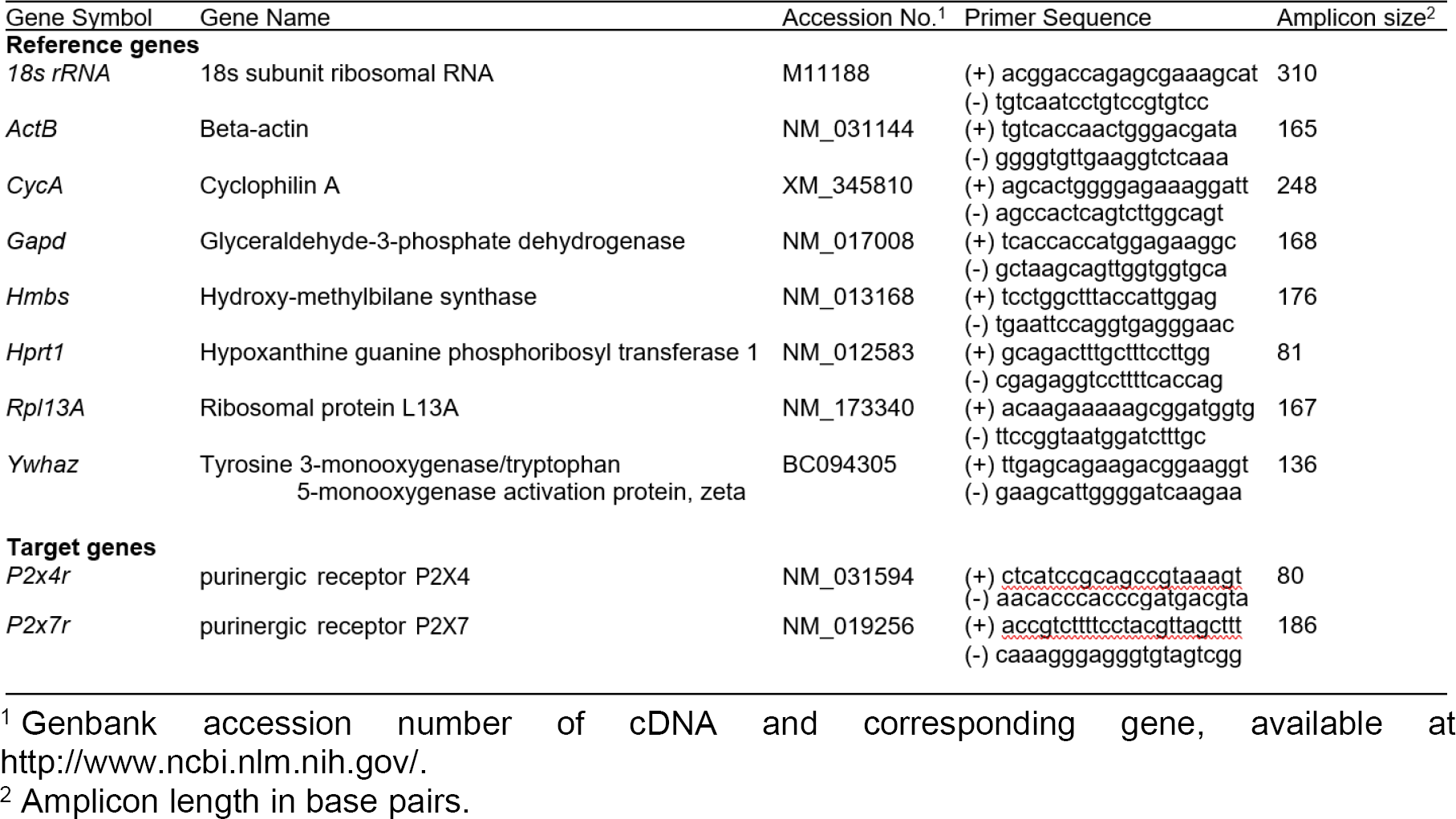
Characteristics of gene-specific real-time qPCR primers (Elfving 2017).

Each SYBR Green reaction (10 µl total volume) contained 1x SYBR Green master mix (Sigma-Aldrich, Denmark), 0.5 µM primer pairs, and 3 µl of diluted cDNA and were carried out as described previously (21). The mixture was heated initially to 95°C for 3 min in to activate hot-start iTaq DNA polymerase and then 40 cycles with denaturation at 95°C for 10 s, annealing at 60°C for 30 s, and extension at 72°C for 60 s were applied. To verify that only one PCR product was detected the samples were subjected to a heat dissociation protocol; after the final cycle of the PCR, the reactions were heat-denatured by increasing the temperature from 60°C to 95°C. The samples and the standard curve were run in duplicate. A standard curve was generated on each plate.

Initially, the mRNA levels were determined for the 8 reference genes. Stability comparison of the expression of the reference genes was then conducted with the Normfinder software. Values of the target genes were subsequently normalized with the geometric mean of the two optimal reference genes (frontal cortex: *Hprt* and *CycA*; ventral hippocampus: *Rpl13A* and *CycA*; dorsal hippocampus: *Hmbs* and *CycA*), based on the NormFinder mathematical algorithm (22).

#### 2.3.2 Western blotting

The concentration (μg/μL) of total proteins was determined in each sample using the Pierce BCA Protein Kit (Thermo Scientific) according to the manufacturer instructions. Tissue samples (prepared using the PARIS purification kit) were mixed with SDS sample buffer (125mM Tris-HCl, pH 6.8; 20% glycerol; 4% SDS; 0.02% bromophenol blue; and 125mM dithiothreitol) (ratio 2:1) and 23 µg of total protein were separated on polyacrylamide gels (Criterion TGX Precast Gel, 26 well comb, 15 μL, 1.0 mm; Bio Rad, #567-1035) and transferred to nitrocellulose membranes (Midi format, 0.2 μm nitrocellulose, single application; Bio Rad, #1704159) using the Trans-Blot Turbo system (Bio Rad, USA; 7 minutes, 25V).

The membranes were washed in TBS (50 mM Tris HCl pH 7.6; 150 mM NaCl) for 5 minutes and incubated with Odyssey Blocking Buffer (OBB; LI-COR Bioscience, #927-5000) for 1 hour at room temperature (RT). Following, they were incubated with primary antibodies to one of the target proteins (rabbit anti-P2×7, 1:200, Alomone Labs, #APR-004; rabbit anti-P2×4, 1:200, Alomone Labs, #APR-002) and to the normalizing protein (mouse anti-β-actin, 1:3000, Licor, #926-42212) overnight at 4°C.

On the following day, the membranes were washed 3 times with 0.1% TBST and incubated with secondary antibody (goat anti-rabbit, 1:10000, Licor, IRDye 800CW; or goat anti-mouse, 1:10000, Licor, IRDye 680RD) diluted in OBB in 0.1% TBST (0.1 % Tween-20 in TBS) (1:2) with 0.01% SDS (to decrease nonspecific background) for 1 hour at RT and protected from light. Still protected from light, the membranes were washed with 0.1% TBST, (3 times, 5 minutes each) and TBS (3 times, 5 minutes each). Infrared signals were detected using the Odyssey CLx infrared imaging system, and bands were quantified using Image Studio software (LI-COR Biosciences). Values from the target proteins were normalized to the β-actin value in each sample.

### 2.4 Data analysis and statistical methods

In the real-time qPCR and WB analysis, respective mRNA levels and normalized fluorescence values were expressed as percentage of control group (FRL rats) and compared by *Student’s t* test. The critical value considered to indicate significant difference between groups was 5% (p<0.05).

## 3 Results and discussion

As illustrated in Figure 1, FSL rats display increased mRNA levels (A: t_12_=2.465, p<0.0297, n=7) of *P2×7r* in FC, but no changes were observed at protein levels (B: t_14_=1.422, p=0, n=8) (Panel 1). Similarly, FSL rats present increased mRNA levels (A: t_12_=6.156, p<0.0001, n=7) of *P2×4r* in FC, with no alterations at protein levels (B: t_14_=0.7788, p=0.4491, n=8) (Panel 2).

**Figure 1.**
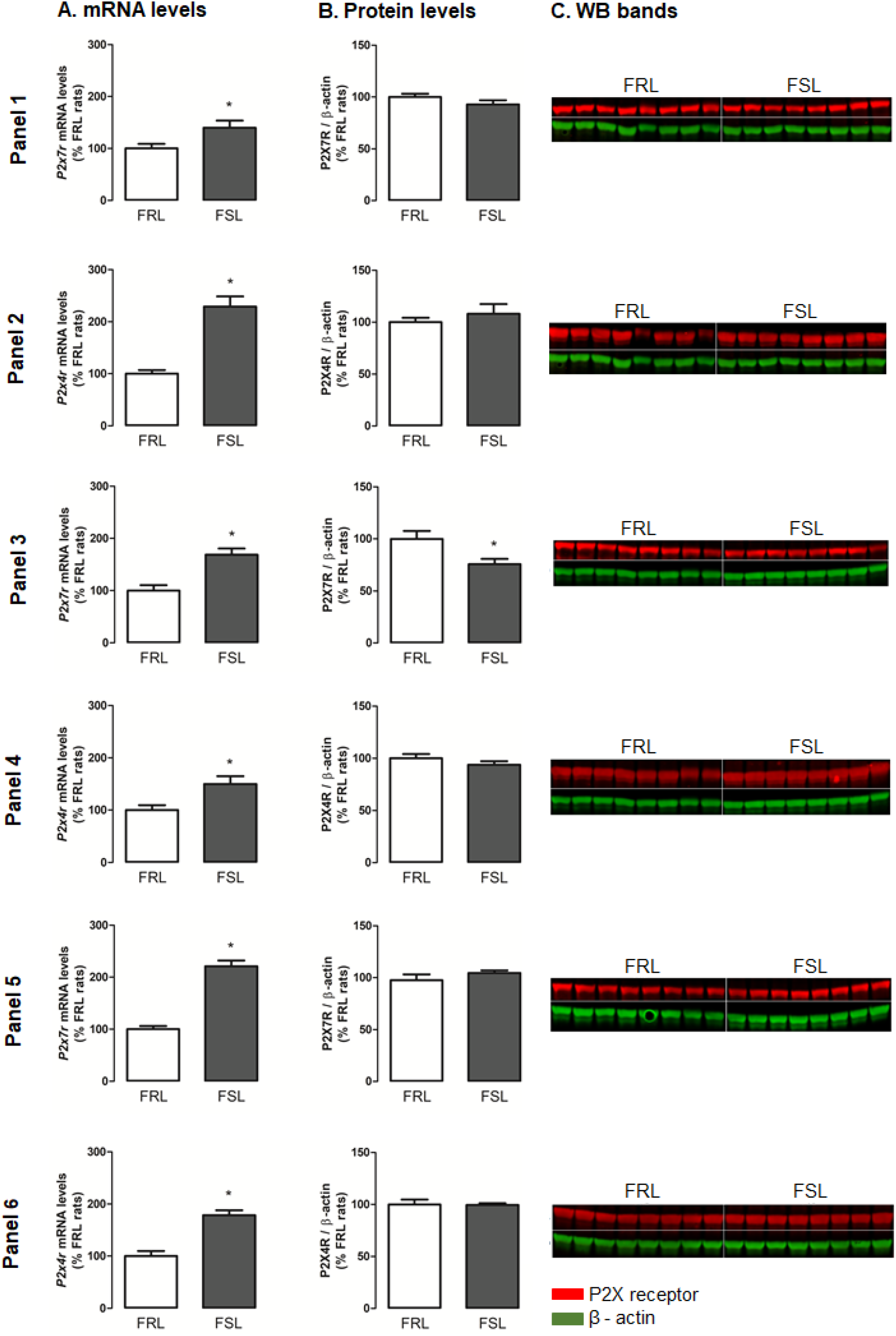
mRNA levels (column A), protein expression (column B) and WB bands (column C) of P2×7R and P2×4R in frontal cortex (Panels 1 and 2), ventral (Panels 3 and 4) and dorsal (Panels 5 and 6) hippocampus of FSL/FRL rats. FSL and FRL rats were decapitated after 1 hour of room habituation and had frontal cortex, ventral and dorsal hippocampus dissected for real-time qPCR and WB analysis. Bars represents mRNA levels of *P2×7r/P2×4r* expressed as percentage of control group (FRL rats) (A) or optical density of these receptors normalized by β-actin also expressed as percentage of FRL rats (B); values are mean ± SEM; asterisks indicate significant differences from FRL rats (*p<0.05), n=7-8 animals/group. WB bands were obtained with Image Studio Lite software (version 5.2) are represented in (C).

FSL rats also show increased mRNA levels (A: t_13_=4.248, p=0.0010, n=7-8) of *P2×7r* in vHip accompanied by a decrease in protein levels (B: t_13_=2.694, p=0.0184, n=7-8) (Panel 3). Panel 4 shows that FSL rats display increased mRNA levels (A: t_13_=2.690, p=0.0186, n=7-8) of *P2×4r* in vHip with no changes on protein levels (B: t_12_=1.124, p=0.2830, n=6-7).

FSL rats showed increased mRNA levels of *P2×7r* in dHip (A: t_14_=9.633, p<0.0001, n=8) with no alterations in protein levels (B: t_14_=1.135, p=0.2756, n=8) (Panel 5). And lastly, FSL rats increased mRNA levels of *P2×4*r in dHip of FSL rats (A: t_14_=5.700, p<0.0001, n=8), with no changes in protein levels (B: t_14_=0.09557, p=0.9252, n=8), as showed in Panel 6.

The main finding in the present study is that FSL rats present increased transcript levels for *P2×7r* and *P2×4r* in FC, ventral and dHip when compared to FRL rats. Despite that, no alterations were observed on protein expression in the analysed structures, except for P2×7R in vHip, which is lower in FSL than in FRL rats.

The activation of the P2×7R (23-31) and P2×4R (32, 33) in the central nervous system is associated to stress response and they can be prevented or reverted by the antidepressant treatment (34-38). However, since no changes were identified at protein levels, the increased mRNA levels can provide an additional level of regulation to stimulate the P2×7R/P2×4R signalling during stress exposure. In line with this suggestion, the mRNA transcripts levels have extremely low predictive value for the expression of their associated proteins, especially in brain samples (39).

Therefore, the functionally dormant transcriptional state of *P2×7r* and *P2×4r* mRNA on FC and Hip of FSL rats could be proposed to be responsible for the increased stress-susceptibility of these animals and the stress exposure could stimulate a rapid synthesis of P2X receptors in FSL rats. Increased levels of these receptors were not observed in the present study, most likely because the animals were not pre-exposed to stress. However, further studies are warranted to further confirm this proposal.

## 4 List of abbreviations

ATP: adenosine triphosphate
dHip: dorsal hippocampus
FC: frontal cortex
FRL: Flinders Resistant Line
FSL: Flinders Sensitive Line
FST: forced swim test
OBB: Odyssey Blocking Buffer
P2×4R: P2×4 receptors
P2×7R: P2×7 receptors
qPCR: quantitative polymerase chain reaction
RT: room temperature
TBS: tris buffered saline
TBST: Tween-20 in TBS
TST: tail suspension test
vHip: ventral hippocampus
WB: Western blotting

## 5 Declarations

### 5.1 Ethics approval and consent to participate

All the procedures were approved by the Danish Animal Experiments Inspectorate (Protocol Number 2012-15-2934-00254). Consent to participate is not applicable to this study since it was conducted in animals.

### 5.2 Consent for publication

Not applicable.

### 5.3 Availability of data and material

The datasets used and/or analysed during the current study are available from the corresponding author on reasonable request.

### 5.6 Authors’ contributions

Deidiane Elisa Ribeiro developed this study as part of her PhD study, she performed the WB and wrote the first version of the manuscript. Heidi Kaastrup Müller helped with the WB analysis. Betina Elfving carried out the real time qPCR experiments. Samia R. L. Joca and Gregers Wegener supervised the study. Gregers Wegener also provided financial and structural support for the development of this project in Aarhus University.

## 5.7 Acknowledgements

Authors are thankful for the invaluable technical assistance of Per Fuglsang Mikkelsen in brain structures dissection, Birgitte Hviid Mumm with the real-time qPCR experiments, and Sanne Nordestgaard Andersen in the WB.

## References

1. Post RM. Transduction of psychosocial stress into the neurobiology of recurrent affective disorder. Am J Psychiatry. 1992;149(8):999–1010.

2. Burnstock G, Krugel U, Abbracchio MP, Illes P. Purinergic signalling: from normal behaviour to pathological brain function. Progress in neurobiology. 2011;95(2):229–74.

3. Alexander SP, Peters JA, Kelly E, Marrion N, Benson HE, Faccenda E, et al. The Concise Guide to PHARMACOLOGY 2015/16: Ligand-gated ion channels. Br J Pharmacol. 2015;172(24):5870–903.

4. Lucae S, Salyakina D, Barden N, Harvey M, Gagne B, Labbe M, et al. P2RX7, a gene coding for a purinergic ligand-gated ion channel, is associated with major depressive disorder. Human molecular genetics. 2006;15(16):2438–45.

5. Nagy G, Ronai Z, Somogyi A, Sasvari-Szekely M, Rahman OA, Mate A, et al. P2RX7 Gln460Arg polymorphism is associated with depression among diabetic patients. Progress in neuro-psychopharmacology & biological psychiatry. 2008;32(8):1884–8.

6. Hejjas K, Szekely A, Domotor E, Halmai Z, Balogh G, Schilling B, et al. Association between depression and the Gln460Arg polymorphism of P2RX7 gene: a dimensional approach. Am J Med Genet B Neuropsychiatr Genet. 2009;150B(2):295–9.

7. Basso AM, Bratcher NA, Harris RR, Jarvis MF, Decker MW, Rueter LE. Behavioral profile of P2×7 receptor knockout mice in animal models of depression and anxiety: relevance for neuropsychiatric disorders. Behav Brain Res. 2009;198(1):83–90.

8. Csolle C, Baranyi M, Zsilla G, Kittel A, Goloncser F, Illes P, et al. Neurochemical Changes in the Mouse Hippocampus Underlying the Antidepressant Effect of Genetic Deletion of P2×7 Receptors. PLoS One. 2013;8(6):e66547.

9. Csolle C, Ando RD, Kittel A, Goloncser F, Baranyi M, Soproni K, et al. The absence of P2×7 receptors (P2rx7) on non-haematopoietic cells leads to selective alteration in mood-related behaviour with dysregulated gene expression and stress reactivity in mice. Int J Neuropsychopharmacol. 2013;16(1):213–33.

10. Pereira VS, Casarotto PC, Hiroaki-Sato VA, Sartim AG, Guimaraes FS, Joca SR. Antidepressant- and anticompulsive-like effects of purinergic receptor blockade: involvement of nitric oxide. Eur Neuropsychopharmacol. 2013;23(12):1769–78.

11. Ma M, Ren Q, Zhang JC, Hashimoto K. Effects of Brilliant Blue G on Serum Tumor Necrosis Factor-alpha Levels and Depression-like Behavior in Mice after Lipopolysaccharide Administration. Clin Psychopharmacol Neurosci. 2014;12(1):31–6.

12. Iwata M, Ota KT, Li XY, Sakaue F, Li N, Dutheil S, et al. Psychological Stress Activates the Inflammasome via Release of Adenosine Triphosphate and Stimulation of the Purinergic Type 2×7 Receptor. Biol Psychiatry. 2016;80(1):12–22.

13. Ribeiro DE, Maiolini VM, Soncini R, Antunes-Rodrigues J, Elias LL, Vilela FC, et al. Inhibition of nitric oxide synthase accentuates endotoxin-induced sickness behavior in mice. Pharmacology, biochemistry, and behavior. 2013;103(3):535–40.

14. Bortolato M, Yardley MM, Khoja S, Godar SC, Asatryan L, Finn DA, et al. Pharmacological insights into the role of P2×4 receptors in behavioural regulation: lessons from ivermectin. The international journal of neuropsychopharmacology / official scientific journal of the Collegium Internationale Neuropsychopharmacologicum. 2013;16(5):1059– 70.

15. Overstreet DH, Wegener G. The flinders sensitive line rat model of depression-25 years and still producing. Pharmacol Rev. 2013;65(1):143–55.

16. Overstreet DH. The Flinders sensitive line rats: a genetic animal model of depression. Neurosci Biobehav Rev. 1993;17(1):51–68.

17. Overstreet DH. Behavioral characteristics of rat lines selected for differential hypothermic responses to cholinergic or serotonergic agonists. Behav Genet. 2002;32(5):335–48.

18. Müller HK, Wegener G, Popoli M, Elfving B. Differential expression of synaptic proteins after chronic restraint stress in rat prefrontal cortex and hippocampus. Brain Res. 2011;1385:26–37.

19. Nava N, Treccani G, Müller HK, Popoli M, Wegener G, Elfving B. The expression of plasticity-related genes in an acute model of stress is modulated by chronic desipramine in a time-dependent manner within medial prefrontal cortex. Eur Neuropsychopharmacol. 2017;27(1):19–28.

20. Bonefeld BE, Elfving B, Wegener G. Reference genes for normalization: a study of rat brain tissue. Synapse. 2008;62(4):302–9.

21. Elfving B, Plougmann PH, Müller HK, Mathe AA, Rosenberg R, Wegener G. Inverse correlation of brain and blood BDNF levels in a genetic rat model of depression. Int J Neuropsychopharmacol. 2010;13(5):563–72.

22. Andersen CL, Jensen JL, Orntoft TF. Normalization of real-time quantitative reverse transcription-PCR data: a model-based variance estimation approach to identify genes suited for normalization, applied to bladder and colon cancer data sets. Cancer Res. 2004;64(15):5245–50.

23. Sperlagh B, Vizi ES. Effect of presynaptic P2 receptor stimulation on transmitter release. Journal of neurochemistry. 1991;56(5):1466–70.

24. Sperlagh B, Kofalvi A, Deuchars J, Atkinson L, Milligan CJ, Buckley NJ, et al. Involvement of P2×7 receptors in the regulation of neurotransmitter release in the rat hippocampus. J Neurochem. 2002;81(6):1196–211.

25. Sperlagh B, Erdelyi F, Szabo G, Vizi ES. Regulation of [H-3]noradrenaline release from the isolated guinea-pig right atrium by P2X receptors located on axon terminals. J Physiol-London. 2000;526:168p-p.

26. Boehm S. ATP stimulates sympathetic transmitter release via presynaptic P2X purinoceptors. Journal of Neuroscience. 1999;19(2):737–46.

27. Witting A, Walter L, Wacker J, Moller T, Stella N. P2×7 receptors control 2-arachidonoylglycerol production by microglial cells. Proceedings of the National Academy of Sciences of the United States of America. 2004;101(9):3214–9.

28. Duan S, Anderson CM, Keung EC, Chen Y, Chen Y, Swanson RA. P2×7 receptor-mediated release of excitatory amino acids from astrocytes. The Journal of neuroscience: the official journal of the Society for Neuroscience. 2003;23(4):1320–8.

29. Wang CM, Chang YY, Kuo JS, Sun SH. Activation of P2X(7) receptors induced [H-3]GABA release from the RBA-2 type-2 astrocyte cell line through a Cl-/HCO3-dependent mechanism. Glia. 2002;37(1):8–18.

30. Walter L, Dinh T, Stella N. ATP induces a rapid and pronounced increase in 2-arachidonoylglycerol production by astrocytes, a response limited by monoacylglycerol lipase. Journal of Neuroscience. 2004;24(37):8068–74.

31. Ballerini P, Rathbone MP, Di Iorio P, Renzetti A, Giuliani P, D’Alimonte I, et al. Rat astroglial P2Z (P2×7) receptors regulate intracellular calcium and purine release. Neuroreport. 1996;7(15-17):2533–7.

32. Guo LH, Trautmann K, Schluesener HJ. Expression of P2×4 receptor by lesional activated microglia during formalin-induced inflammatory pain. Journal of neuroimmunology. 2005;163(1-2):120–7.

33. Schwab JM, Guo L, Schluesener HJ. Spinal cord injury induces early and persistent lesional P2×4 receptor expression. Journal of neuroimmunology. 2005;163(1-2):185–9.

34. Bliss TV, Cooke SF. Long-term potentiation and long-term depression: a clinical perspective. Clinics. 2011;66 Suppl 1:3–17.

35. Kreisel T, Frank MG, Licht T, Reshef R, Ben-Menachem-Zidon O, Baratta MV, et al. Dynamic microglial alterations underlie stress-induced depressive-like behavior and suppressed neurogenesis. Molecular psychiatry. 2014;19(6):699–709.

36. Pariante CM, Lightman SL. The HPA axis in major depression: classical theories and new developments. Trends in neurosciences. 2008;31(9):464–8.

37. Veith RC, Lewis N, Linares OA, Barnes RF, Raskind MA, Villacres EC, et al. Sympathetic nervous system activity in major depression. Basal and desipramine-induced alterations in plasma norepinephrine kinetics. Arch Gen Psychiatry. 1994;51(5):411–22.

38. Barton DA, Dawood T, Lambert EA, Esler MD, Haikerwal D, Brenchley C, et al. Sympathetic activity in major depressive disorder: identifying those at increased cardiac risk? Journal of hypertension. 2007;25(10):2117–24.

39. Bauernfeind AL, Babbitt CC. The predictive nature of transcript expression levels on protein expression in adult human brain. BMC Genomics. 2017;18(1):322.

